# Serotonergic neuron ribosomes regulate the neuroendocrine control of *Drosophila* development

**DOI:** 10.1101/2021.06.10.447971

**Authors:** Lisa P. Deliu, Deeshpaul Jadir, Abhishek Ghosh, Savraj S. Grewal

## Abstract

The regulation of ribosome function is a conserved mechanism of growth control. While studies in single cell systems have defined how ribosomes contribute to cell growth, the mechanisms that link ribosome function to organismal growth are less clear. Here we explore this issue using *Drosophila Minutes*, a class of heterozygous mutants for ribosomal proteins (Rps). These animals exhibit a delay in larval development caused by decreased production of the steroid hormone ecdysone, the main regulator of larval maturation. We found that this developmental delay is not caused by decreases in either global ribosome numbers or translation rates. Instead, we show that they are due in part to loss of Rp function specifically in a subset of serotonin (5-HT) neurons that innervate the prothoracic gland to control ecdysone production. We found that these 5-HT neurons have defective secretion in *Minute* animals, and that overexpression of synaptic vesicle proteins in 5-HTergic cells can partially reverse the *Minute* developmental delay. These results identify a cell-specific role for ribosomal function in the neuroendocrine control of animal growth and development.

## INTRODUCTION

The regulation of ribosome and protein synthesis are conserved mechanisms of growth control. Several decades of studies in unicellular systems such as E. coli, yeast and cultured mammalian cells have defined both the signaling pathways that couple growth cues to ribosome synthesis and function, and the mechanisms by which changes in mRNA translation drive cell growth and proliferation (Dai and Zhu, 2020; Lempiainen and Shore, 2009; Rudra and Warner, 2004; Warner, 1999). However, the mechanisms that operate in whole animals during developmental growth are less clear. In these contexts, body growth is not determined solely by processes that govern cell-autonomous growth, but also by inter-organ communication to ensure coordinated growth and development across all tissues and organs (Boulan et al., 2015; Droujinine and Perrimon, 2016; Grewal, 2012). Hence, tissue specific changes in ribosome function have the potential to mediate non-autonomous effects on whole-body physiology to control organismal development.

The complex links between ribosome function and animal development are exemplified by the organismal biology of ribosomal proteins (Rps) (Terzian and Box, 2013; Xue and Barna, 2012). Metazoan ribosomes have 70-80 Rps, and mutants for almost all of these are homozygous lethal in animals, emphasising their essential role in ribosome synthesis and function. However, in many cases Rp mutants show dominant phenotypes as heterozygotes. These phenotypes are often specific to the affected Rp and can give rise to tissue-specific effects that cannot be explained simply by lowered overall protein synthesis and growth rates. For example, in zebrafish certain *rp/+* mutants can develop peripheral nerve tumors (Amsterdam et al., 2004). Similarly, some *Drosophila* Rp mutants develop selective tissue overgrowth phenotypes (Torok et al., 1999; Watson et al., 1992). Several *Rp/+* mutants in mice have also been shown to each exhibit tissue specific developmental defects that differ based on the Rp affected. For example, *rpl38/+* mice show specific skeletal segmentation defects (Kondrashov et al., 2011), *rps14/+* mice show defects in blood development (Barlow et al., 2010), and *rpl27a/+ mice* show defects in cerebellar development (Terzian et al., 2011). The dominant effects of *rp/+* mutations also extend to humans, where several pathologies, collectively termed ribosomopathies, are caused by heterozygosity for Rp mutations, and lead to tissue-specific effects such as blood disorders, congenital growth defects, and predisposition to cancer (Farley-Barnes et al., 2019; Kampen et al., 2020; Yelick and Trainor, 2015). The mechanisms that determine these dominant effects of *rp/+* mutations are not fully clear but are thought to involve selective alterations in mRNA translation. These alterations may occur either as a result of lowered ribosome numbers or due to ribosome heterogeneity, where ribosomes with different complements of Rps have been proposed to have different translational properties (Dinman, 2016; Genuth and Barna, 2018a, b; Khajuria et al., 2018; Mills and Green, 2017). These studies emphasise the importance of further work to understand how Rp function contributes to organismal growth and development.

*Drosophila* larvae have provided an excellent model system in which to define the cell-, tissue- and body-level mechanisms that control developmental growth (Andersen et al., 2013; Boulan et al., 2015; Texada et al., 2020). Larvae grow almost 200-fold in mass over 4-5 days before undergoing metamorphosis to the pupal stage. This developmental transition is controlled by a pulse of secretion of the steroid hormone, ecdysone, from the prothoracic gland (PG), which then acts on tissues to stimulate pupation at the end of the larval period (Kannangara et al., 2021; Pan et al., 2021; Yamanaka et al., 2013). The timing of this pulse is under control of two separate subsets of neurons expressing either the neuropeptide, PTTH, or the neuromodulator serotonin (5-HT), that each innervate the PG and stimulate ecdysone production (McBrayer et al., 2007; Shimada-Niwa and Niwa, 2014; Shimell et al., 2018). This neuroendocrine network integrates signals from the environment and other tissues to ensure proper timing of the ecdysone pulse and the larval-pupal transition. For example, nutrient signals can act on both the 5-HT neurons and the PG to ensure proper coupling of development maturation with nutrients (Layalle et al., 2008; Shimada-Niwa and Niwa, 2014). Epithelial disc damage also leads to a delay in larval development to allow time for proper tissue regeneration before transition to the pupal stage. One way that this delay is mediated is by suppression of PTTH signaling by dilp8, an insulin/relaxin-like peptide that signals from damaged discs to a subset of Lgr3 receptor expressing neurons that inhibit PTTH neuronal activity (Colombani et al., 2015; Colombani et al., 2012; Garelli et al., 2012; Garelli et al., 2015; Jaszczak et al., 2016). In addition, the inflammatory cytokine, Upd3 can signal directly from damaged discs to the PG to suppress ecdysone and delay development (Romao et al., 2021).

An interesting class of mutants that exhibit alterations in larval development are the *Minutes* (Lambertsson, 1998; Marygold et al., 2007). These are dominant mutants that are classically described by their developmental delay and short bristles. Almost all *Minutes* are *rp/+* mutants and they have perhaps been best studied in context of cell competition, a process in which mosaic clones of *rp/+* cells in imaginal disc epithelia are outcompeted and killed by surrounding wild-type (*+/+*) cells. Several mechanisms have been described to account why *rp/+* cells are outcompeted including altered proteostasis (Baumgartner et al., 2021; Recasens-Alvarez et al., 2021), competition for *dpp* growth factor (Moreno et al., 2002), induction of innate immune signaling (Germani et al., 2018; Meyer et al., 2014), and induction of the transcription factor Xrp1 (Baillon et al., 2018; Lee et al., 2018). Interestingly, some of these disc-intrinsic, cell competition effects have also been shown to partially account for the organismal delay in development seen in *rp/+* animals. For example, disc-specific Rp knockdown stimulates Xrp1 induction of dilp8 (Boulan et al., 2019), and both loss of Xrp1 and disc-specific knockdown of dilp8 can each partially reverse the delay in development seen in *rp/+* animals (Akai et al., 2021; Ji et al., 2019; Lee et al., 2018). It has also been shown that loss of Rp function specifically in the PG can also explain the overall delay in development in *rps6/+* animals (Lin et al., 2011). These results suggest that the overall delay in organismal development seen in *rp/+* animals may result from tissue non-autonomous effects of Rps. Hence *Minutes* provide an excellent system to explore how tissue-selective functions of Rps contribute to whole-body phenotypes.

Here we provide further evidence for tissue-specific effects of Rp in the control of larval development. We describe how loss of Rp function results in defects in vesicle-mediated secretion specifically in the 5-HT neurons that innervate the PG leading to developmental delay in *Minute* animals.

## RESULTS

### Rps13/+ animals do not show a global decrease in ribosome levels or protein synthesis

For our study we used flies heterozygous for previously characterized allele of ribosomal protein S13, *P[lacW]M (2)32A* (hereafter referred to as *rpS13/+* animals), which have decreased expression of *rps13* mRNA (Figure S1A) and have been observed to have the classic *Minute* phenotype of shorter and thinner bristles and a delay in larval development (Saeboe-Larssen et al., 1998). We quantified the delay in development of *rpS13/+* and controls (*w^1118^*) by measuring the time it took for animals to reach the pupal stage after egg laying. We found that *rpS13/+* animals were delayed in development by about 40 hours, which corresponds to a delay of approximately 20% compared to control animals (Figure 1A). We also measured body size as the larvae developed, and we saw that *rpS13/+* larvae had a smaller size compared to age-matched control animals at different stages of larval development (Figure S1B). However, due to their prolonged larval period, the *rpS13/+* animals grew for a longer time. Hence, when we measured both final larval and pupal size, we found that, in both cases, *rpS13/+* animals were about 12% larger than controls (Figure 1B, C, Figure S1C). We measured mouth hook movements as measure of feeding rate and saw a small, but significant, increase in *rpS13/+* larvae when compared to controls (Figure S1D). This indicates that the growth and developmental phenotypes of *rpS13/+* animals do not result simply from reduced feeding. These data suggest that *rpS13/+* animals exhibit a reduced growth and developmental rate.

**Figure 1.**
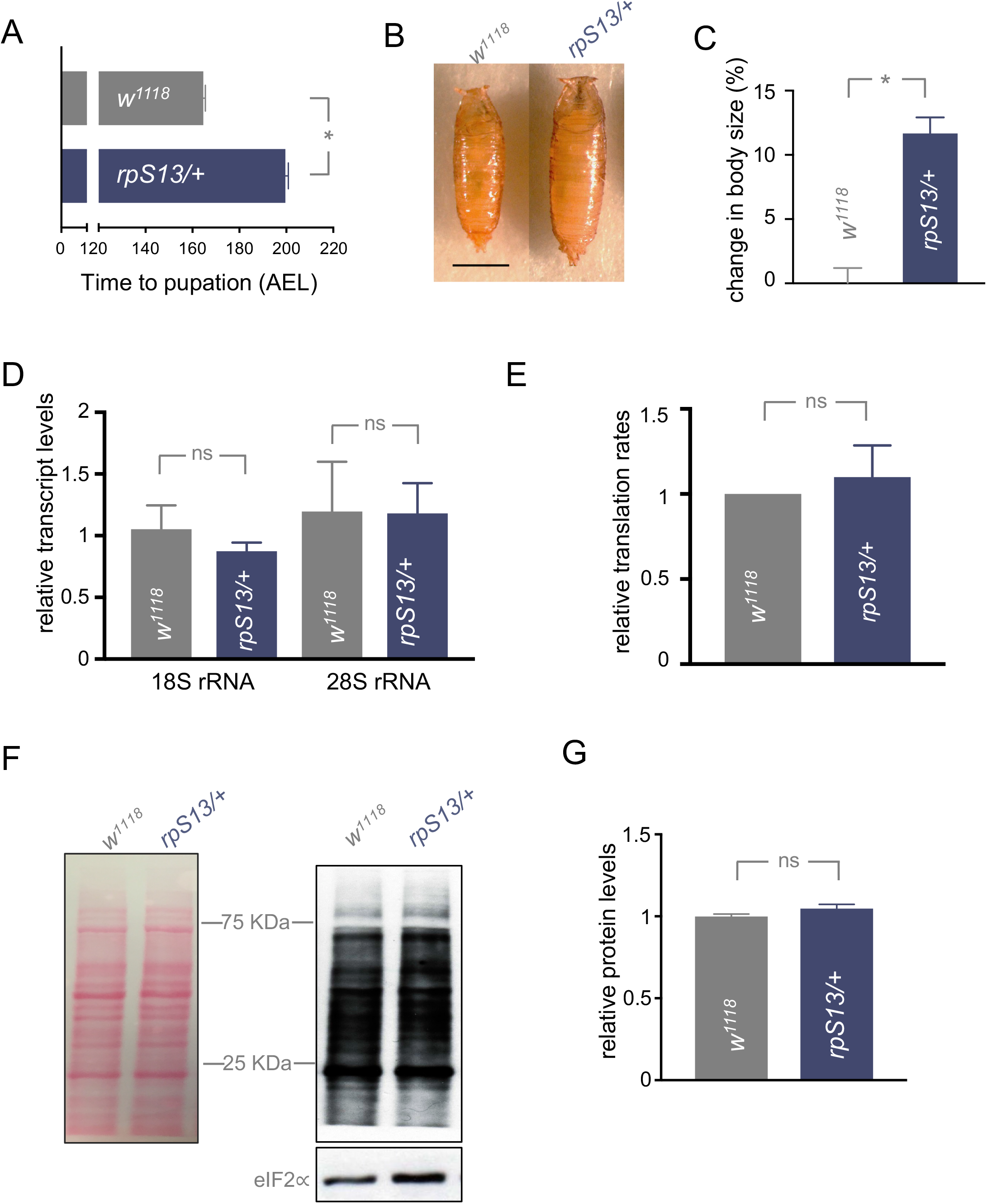
Developmental delay of *rpS13/+* animals is not due to a decrease in ribosome numbers or reduced translation rates. (A) Developmental timing from larval hatching to pupation of *w^1118^* and *rpsS13/+* animals, n=147 and 170 respectively. *rpS13/+* animals are on average 35 hours delayed when compared to their wild-type controls. Data are presented as +/- SEM. *p < 0.05, Mann-Whitney U test. (B) Representative images of *W^1118^* and *rpS13/+* pupae. Scale bar, 1mm. (C) Change in pupal volume of *rpS13/+* (n = 212) when compared to *w^1118^* controls (n = 229). *rpS13/+* animals grow on average 11% larger than controls. Data are presented as +/- SEM. *p < 0.05, Mann-Whitney U test. (D) Transcript levels of 18S and 28S rRNA in *w^1118^* and *rpsS13/+* show no reduction in ribosomal rRNA in *rpsS13/+* heterozygotes. mRNA was isolated from third instar wandering larvae. Total RNA was isolated and measured by qRT-PCR, n = 4 independent samples per genotype. Data are presented as +/- SEM. p > 0.05, Student’s t-test. (E) Relative translation rates based on quantification of puromycin staining, n = 6 independent samples per genotype. Data are presented as +/- SEM. p > 0.05, Student’s t-test. (F) Puromycin labelling assay shows no difference in *rpS13/+* translational rates in third instar wandering larvae when compared to controls. Left, Ponceau S staining showing total protein. Right, anti - puromycin and anti - eIF2∝ (loading control) immunoblots. (G) Relative protein concentration levels from third instar wandering *w^1118^* and *rpsS13/+* larvae. Absorbance was measured at 465nm using the Bradford assay, n =5 independent samples per genotype. Data are presented as +/- SEM. p > 0.05, Student’s t-test.

Studies in different model systems have shown that the phenotypes seen in rp/+ animals are often associated with lowered ribosome numbers and reduced protein synthesis. We therefore investigated ribosome levels and protein synthesis in *rpS13/+* animals. In order to measure ribosome numbers, we measured mature 18S and 28S rRNA in wandering L3 whole larval lysates. We saw no significant difference in rRNA levels between *rpS13/+* and control larvae (Figure 1D). Total protein content in wandering L3 larval lysates also showed no significant difference in *rpS13/+* larvae compared to control larvae (Figure 1E). Finally, we investigated whether *rpS13/+* animals show a decrease in protein synthesis rate. To do this we used a puromycin labelling assay (Deliu et al., 2017). We first quantified the levels of puromycin incorporation of *rpS13/+* and control animals at the wandering larval stage in order to developmentally match control and *rpS13/+* animals and found no significant difference in protein synthesis rates (Figure 1F, G). We repeated this assay at two other earlier time points with aged-matched larvae, and once again found no decrease in translation rates in *rpS13* /+ larvae compared to control larvae (Figure S2). This suggests that the *Minute* delayed development is not due to a global loss of ribosome numbers or translational capacity.

### *rpS13/+* animals show a defect in ecdysone signalling

The duration of the larval period is controlled in large part by the steroid hormone, ecdysone (Pan et al., 2021). In particular, at the end of larval development, a neuro-endocrine circuit stimulates a pulse of ecdysone production and secretion from the prothoracic gland (PG). This circulating ecdysone then acts on larval tissues to trigger the larval to pupal transition. Any defects in this neuro-endocrine circuit leads to a delay in larval development to the pupal stage. Given their delayed development, we examined whether *rpS13/+* animals show a defect in ecdysone signaling. We did this by measuring the transcript levels of *phantom* and *spookier* both of which encode enzymes for PG ecdysone production. As previously described, both showed maximal expression peaks at 120 hours AEL in control animals, consistent with the ecdysone pulse that triggers pupation (Figure 2A). However, in *rpS13/+* animals these peaks were delayed by about one day (144 hours) and continued to show expression even at 168 hours for larvae that were still wandering (Figure 2A), suggesting a delay in ecdysone signalling. We also found that feeding larvae 20 hydroxyecdysone (20HE) was able to partially reverse the development timing delay seen in the *rpS13/+* by about one third of the total delay (Figure 2B). Ecdysone synthesis in the PG can be stimulated by several different signaling pathways, including the Ras/ERK and TOR kinase pathways (Cruz et al., 2020; Layalle et al., 2008; Rewitz et al., 2009). When we overexpressed the TOR activator, Rheb in the PG, we found that while it had no effect on developmental timing in control animals, it was sufficient to partially reverse the development timing delay seen in the *rpS13/+* animals again by about one third of the total delay (Figure 2C). Together, these data indicate that *rpS13/+* animals exhibit a delay in development that can be explained in part due to blunted ecdysone signaling.

**Figure 2.**
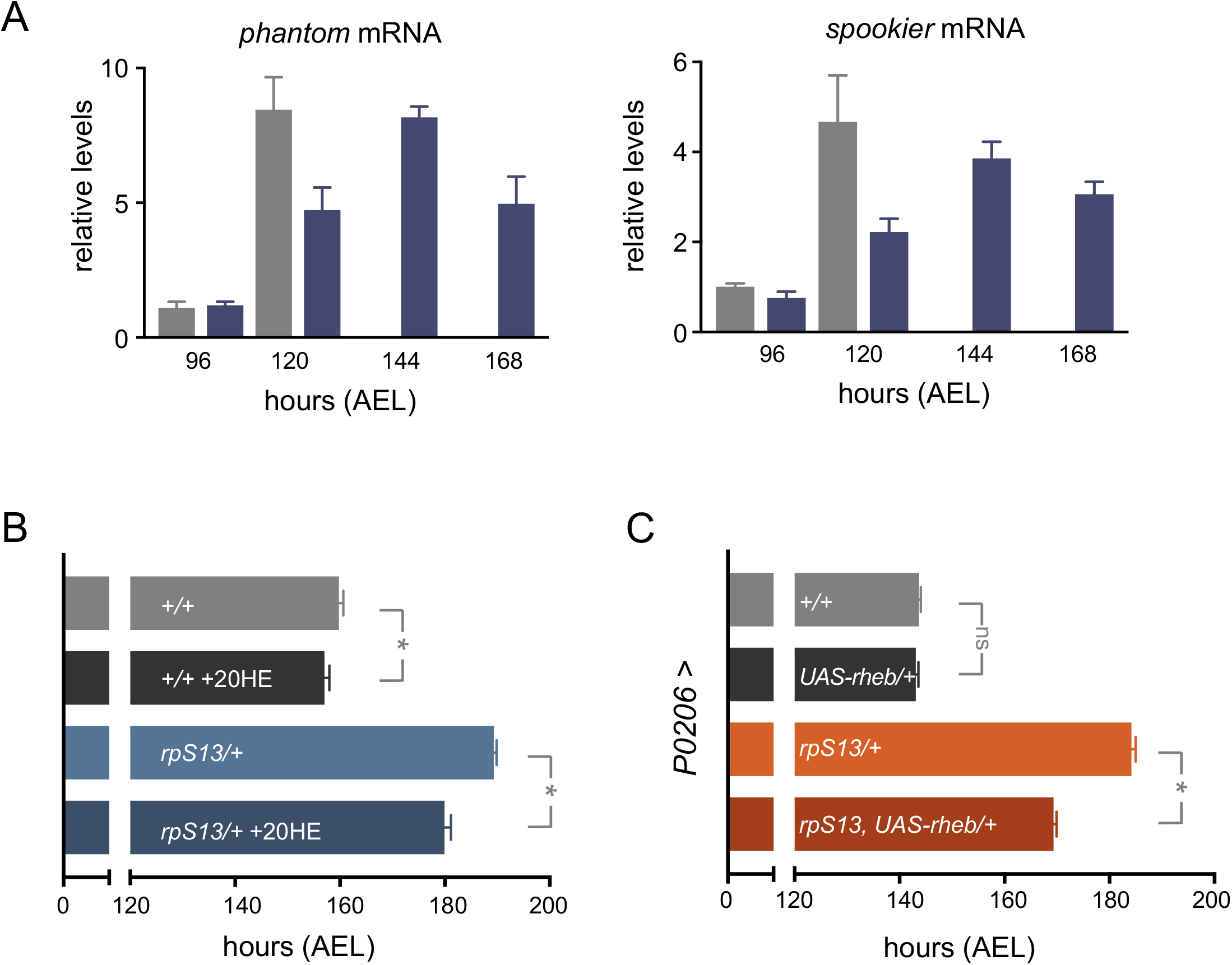
Delayed development in *rpS13/+* animals is due to impaired ecdysone production. (A) Transcript levels of two *Halloween* genes, *phantom* and *spookier* expression are delayed in *rpS13/+* larvae when compared to wild-type controls. mRNA was isolated from third instar larvae every 24 hours beginning with the 96 hours AEL time point until pupation of each respective genotype. Total RNA was isolated and measured by qRT-PCR, n = 4 independent samples per genotype. Data are presented as +/- SEM. (B) Ecdysone (20HE) was supplemented into food at a 0.3mg/mL concentration in both *rpS13/+* and *w^1118^* animals, n = 92, n = 116, respectively. 20HE partially rescued *rpS13/+* developmental delay by 10 hours (~33 %). Controls for both *w^1118^* and *rpS13/+* were fed the same concentration (0.3mg/mL) of 95% ethanol in food, n = 142, n = 148, respectively. Data are presented as +/- SEM. *p < 0.05, Mann-Whitney U test. (C) *UAS-rheb* expression in the prothoracic gland using *P0206-Gal4* partially rescues the developmental delay of *rpS13/+* larvae by 15 hours (~36 %). No difference is seen between *+/+* controls and *UAS-rpS13* overexpression. *+/+* n = 152, *UAS-rheb/+* n = 157, *rpS13/+* n = 151, *UAS-rheb/rpS13* n = 144. Data are presented as +/- SEM. *p < 0.05, Mann-Whitney U test.

### 5-HT neuronal *rpS13* is required for proper developmental timing

Our earlier data indicated that *rpS13/+* animals did not show any global changes in whole-body ribosome or protein synthesis levels. Hence, it is possible that the delayed development in *rpS13/+* animals reflects a more selective role for RpS13, perhaps in specific cells or tissues involved in controlling ecdysone. To investigate this possibility, our approach was to use the *Gal4/UAS* system to re-express a RpS13 transgene (*UAS-rpS13*) in specific tissues in *rpS13/+* animals and then examine whether this could reverse the delay in development. We focused in particular on examining cells and tissues important for stimulating the ecdysone pulse. We first re-expressed *UAS-rpS13* the PG using the PG driver, *P0206-Gal4*. We saw no significant change in timing in control animals with the driver alone or with the over-expression of *UAS-rpS13* in control animals. Moreover, we found that PG-specific expression of *UAS-rpS13* in *rpS13/+* larvae was unable to reverse the delay in development seen in the *rpS13/+* animals (Figure 3A). We then examined the imaginal discs. Growth aberrations caused by reduced Rp expression lead to release of dilp8 which ultimately has a negative effect on ecdysone synthesis. We used the *esg-Gal4^ts^* which directs temperature-inducible expression in all imaginal cells in the larvae. We found that expression of *UAS-rpS13* throughout larval development using *esg-Gal4^ts^* had little effect on developmental timing in control animals, but instead had an exacerbation of developmental delay in *rpS13/+* animals (Figure 3B). The PG-induced expression of ecdysone at the end of the larval period is controlled by neuronal signals to the PG. We therefore examined the effect of re-expressing *rpS13* in neurons using a pan-neuronal driver, *elav-Gal4*. We found that while neuronal expression of *UAS-rpS13* in the control animals did not affect timing, it was sufficient to partially rescue the developmental delay in *rpS13/+* larvae. Indeed, this rescue was similar to that seen with either 20HE feeding or by stimulation of TOR signaling in the PG (Figure 3C). We confirmed this result by using a second pan-neuronal driver, *nSyb-Gal4*, which also rescued timing by roughly one third while not accelerating timing in the wild type animals (Figure 3D). Since *rpS13/+* animals also have an increased final body size phenotype, we measured pupal volume in animals with neuronal *UAS-rpS13* expression. We found that expression of *UAS-rpS13* in the *rpS13/+* larvae with either *elav-Gal4* or *nsyb-Gal4* led to a significant reversal of the increased body size seen in *rpS13/+* animals (Figure 3E, F). These data point to a neuronal requirement for RpS13 in larval development that accounts for the ecdysone defect in *Minute* animals.

**Figure 3.**
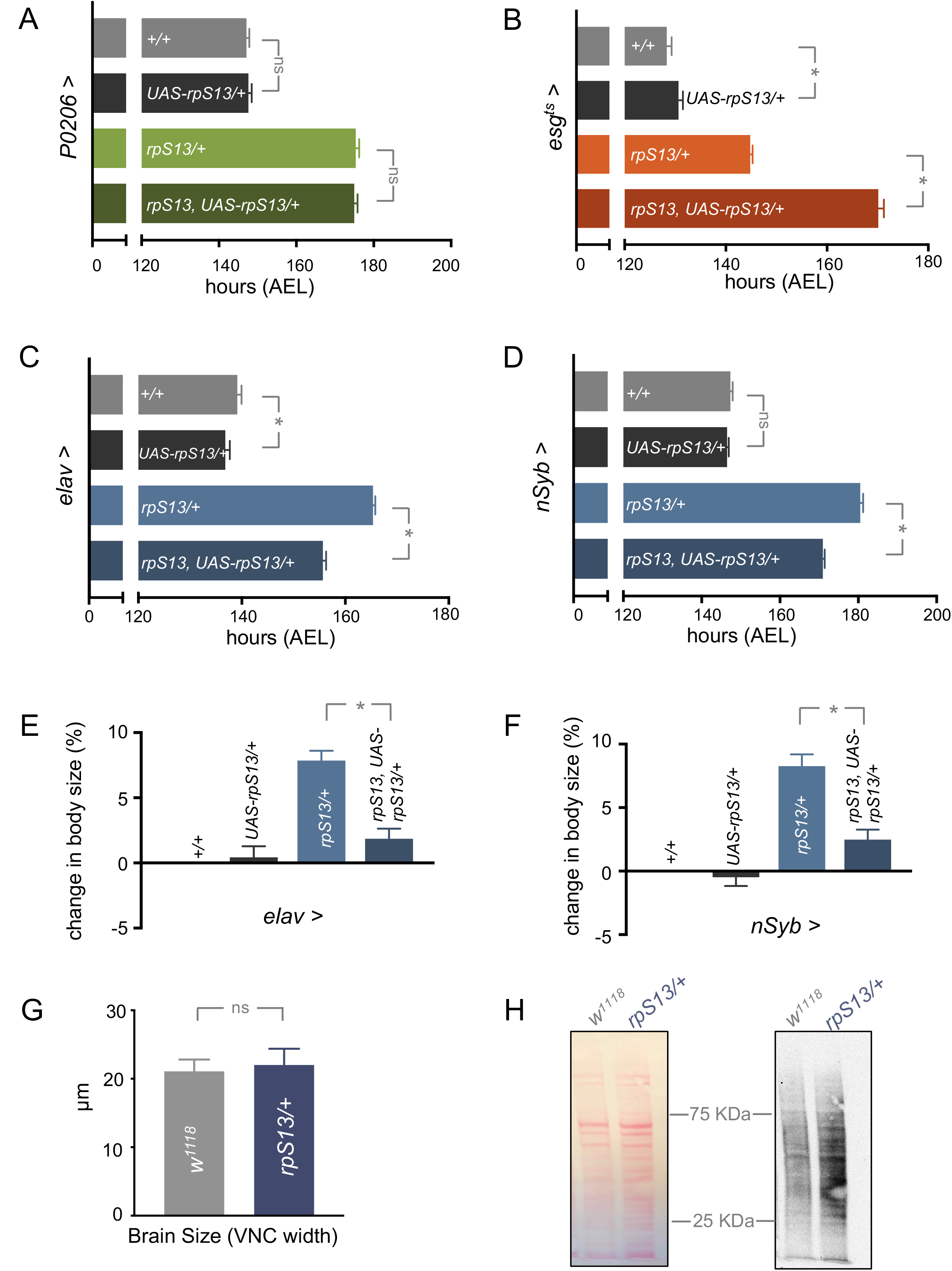
*rpS13* re-expression in neurons partially reverses the *rpS13/+* developmental delay. (A) *rpS13* re-expression in the prothoracic gland using *P0206-Gal4* does not rescue the developmental delay of *rpS13/+* larvae. No difference is seen between *+/+* controls (n = 95) and *UAS-rpS13* overexpression (n = 95) or between *rpS13/+* (n = 98) and *rpS13, UAS-rpS13/+* (n = 87) animals. Data are presented as +/- SEM. p > 0.05, Mann-Whitney U test. (B) *rpS13* re-expression in the imaginal discs using *esg^ts^-Gal4* does not rescue the developmental delay of *rpS13/+* larvae. There is a minor increase in developmental timing between *+/+* controls (n = 96) and *UAS-rpS13* overexpression (n = 29) and an enhanced delay between *rpS13/+* (n = 99) and *rpS13, UAS-rpS13/+* (n = 86) animals. Data are presented as +/- SEM. *p < 0.5, Mann-Whitney U test. (C) Pan-neuronal expression of *rpS13* in *rpS13/+* animals using *elav-Gal4* partially rescues the developmental delay by 10 hours (~33 %). *+/+* n = 180, *rpS13/+* n = 167, *UAS-rpS13/+* n = 182, *rpS13, UAS-rpS13/+* n = 226. Data are presented as +/- SEM. *p < 0.5, Mann-Whitney U test. (D) Expression of *rpS13* using a second pan-neuronal driver, *nSyb-Gal4* also partially rescues *rpS13/+* developmental delay by 10 hours (~33 %). *+/+* n = 182, *rpS13/+* n = 180, *UAS-rpS13/+* n = 190, *rpS13, UAS-rpS13/+* n = 176. Data are presented as +/- SEM. *p < 0.05, Mann-Whitney U test. (E) Pan-neuronal expression of *rpS13* in *rpS13/+* animals using *elav-Gal4* partially restores *rps13/+* overgrowth phenotype. *+/+* n = 313, *rpS13/+* n = 468, *UAS-rpS13/+* n = 456, *rpS13, UAS-rpS13/+* n = 475.Data are presented as +/- SEM. *p < 0.05, Mann-Whitney U test. (F) Pan-neuronal expression of *rpS13* in *rpS13/+* animals using *nSyb-Gal4* partially restores *rps13/+* overgrowth phenotype. *+/+* n = 588, *rpS13/+* n = 484, *UAS-rpS13/+* n = 586, *rpS13, UAS-rpS13/+* n = 536.Data are presented as +/- SEM. *p < 0.05, Mann-Whitney U test. (G) Ventral nerve cord (VNC) width (μm) was used to compare brain sizes in third instar, wandering larvae of *+/+* controls (n = 12) and *rpS13/+* (n = 12) animals. Data are presented as +/- SEM. p > 0.05, Student’s t-test. (H) Puromycin labelling assay shows no decrease, but rather a small increase in *rpS13/+* translation rates in third instar wandering larvae brains when compared to controls, *+/+*. Left, Ponceau S staining showing total protein. Right, anti-puromycin immunoblot.

We examined whether we could see any global changes in either the size or protein synthesis levels of brains from *rpS13/+* animals compared to controls. However, when we measured wandering larval brain size (ventral nerve cord width) or translation rates (using puromycin labelling) and found no decrease in the *Minute* animals (Figures 3G, H). We therefore examined whether the requirement for neuronal RpS13 for proper developmental timing might reflect a role in a specific subset of neurons, in particular those known to influence PG function. One important subset is a pair of bilateral PTTH-expressing neurons that directly innervate the PG. These respond to developmental cues to secrete the peptide PTTH which acts on the PG to stimulate peak levels of ecdysone biosynthesis at the end of the larval stage (McBrayer et al., 2007; Shimell et al., 2018). The PTTH neurons are also themselves directly regulated by another subset of neurons (Lgr3-expressing neurons) that are controlled by tissue damage (Colombani et al., 2015; Garelli et al., 2015; Jaszczak et al., 2016). We therefore examined the effects of expression of *UAS-rpS13* in these neurons using the *ptth-Gal4* and *lgr3-Gal4* drivers. We found that when we expressed *UAS-rpS13* in PTTH neurons we saw no effect on developmental timing in control animals and a small rescue of the developmental delay in *rps13/+* animals (Figure 4A). Expression of *UAS-rpS13* using the *lgr3-Gal4* driver had no effect on developmental timing either in control or *rps13/+* animals (Figure 4B). These results suggest that expression of the RpS13 in neurons that control PTTH signaling does not fully account for the rescue of *Minute* developmental timing that we observed with pan-neuronal RpS13 expression.

**Figure 4.**
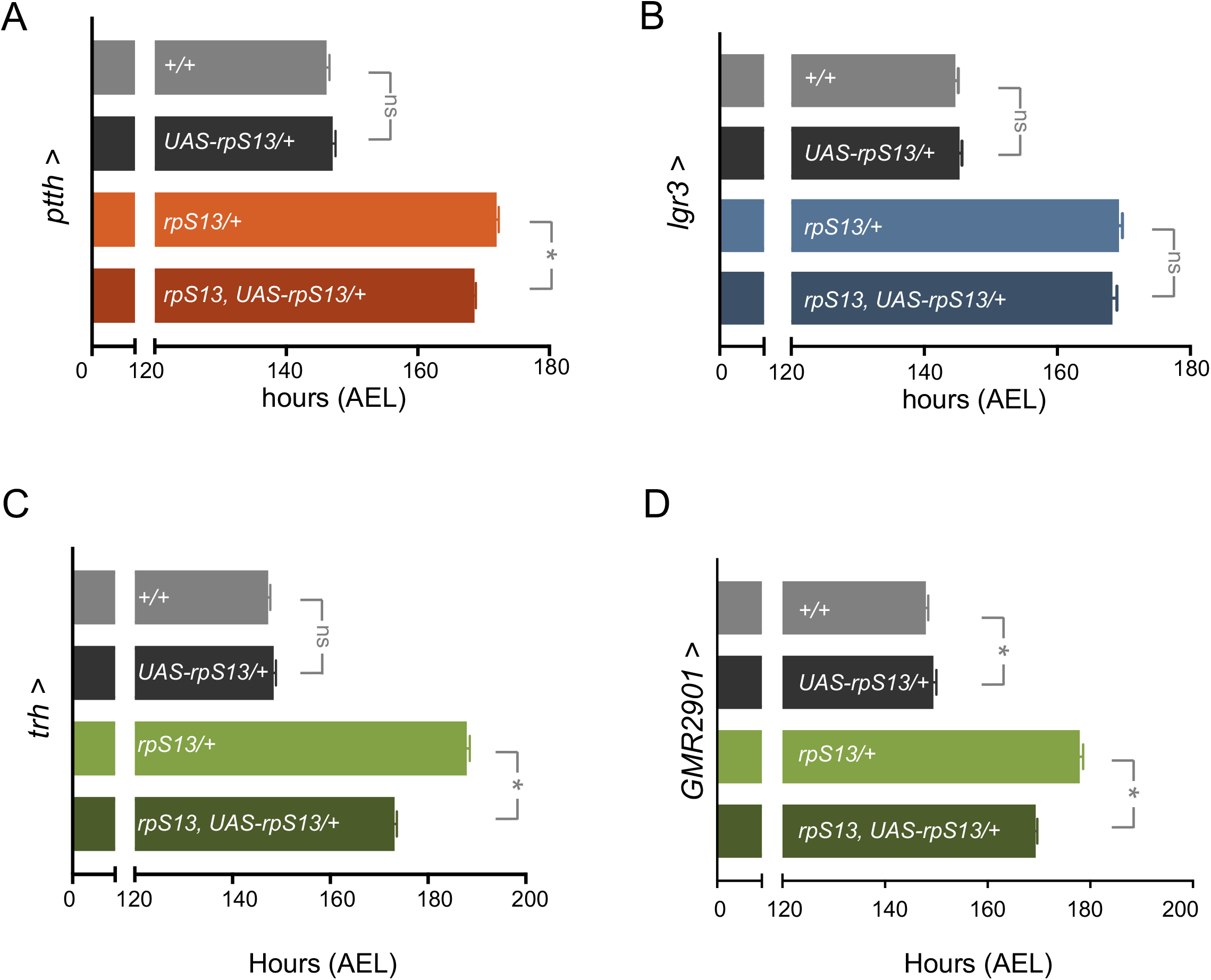
*rpS13* re-expression in the serotonergic neurons that innervate the prothoracic gland partially rescues *rpS13/+* developmental delay. (A) *rpS13* expression in the ptth neurons using *ptth-Gal4* has a very mild effect on the developmental delay of *rpS13/+* larvae. No difference is seen between *+/+* controls (n = 233) and *UAS-rpS13* overexpression (n = 288) or between *rpS13/+* (n = 282) and *rpS13, UAS-rpS13/+* (n = 282) animals. Data are presented as +/- SEM. *p < 0.05, Mann-Whitney U test. (B) *rpS13* expression in the Lgr3 neurons using *Lgr-Gal* has no effect on developmental timing of *rpS13/+* or control animals. *+/+* n = 192 *rpS13/+* n = 176, *UAS-rpS13/+* n = 177, *rpS13, UAS-rpS13/+* n = 166. Data are presented as +/- SEM. *p < 0.05, Mann-Whitney U test. (C) *rpS13* expression in the serotonergic neurons using *Trh-Gal4* partially rescues developmental delay of *rpS13/+* larvae by 14 hours (~36 %). *+/+* n = 241 *rpS13/+* n = 176, *UAS-rpS13/+* n = 234, *rpS13, UAS-rpS13/+* n = 230. Data are presented as +/- SEM. *p < 0.05, Mann-Whitney U test. (D) *rpS13* expression in a subset of serotonergic neurons that directly innervate the prothoracic gland, using *GMR29H01-Gal4* partially rescues developmental delay of *rpS13/+* larvae by 9 hours (~30 %). *+/+* n = 225 *rpS13/+* n = 196, *UAS-rpS13/+* n = 192, *rpS13, UAS-rpS13/+* n = 248. Data are presented as +/- SEM. *p < 0.05, Mann-Whitney U test.

The PG is also directly innervated by serotonergic (5-HT) neurons (Shimada-Niwa and Niwa, 2014). These 5-HT neurons are required for proper ecdysone production at the end of the larval stage, particularly in response to dietary nutrients. When we used the 5-HT neuronal driver *trh-Gal4* to express *UAS-rpS13* we found that we could reverse the delay in development seen in *rpS13/+* animals by about one-third. This recapitulates the extent of the rescue seen with pan-neuronal *UAS-rpS13* expression and is similar to that seen with either 20HE feeding or by TOR-dependent activation in the PG (Figure 4C). There are approximately 100 serotoninergic neurons in the larval brain, of which three pairs innervate the prothoracic gland directly (Shimada-Niwa and Niwa, 2014). Using another neuronal driver which has a more limited expression pattern which includes these three pairs of 5-HT neurons we re-expressed *UAS-rpS13* in *rpS13/+* and control animals and found that timing was again rescued by approximately one-third in the *rpS13/+* animals while development was unaffected in control animals (Figure 4D). These data suggest a specific role for RpS13 in 5-HT neurons that innervate the PG in the regulation of developmental timing.

We speculated that reduced RpS13 levels in 5-HT neurons may lead to decreased protein synthesis, thus we examined the effects of expressing two known stimulators of translation, *dMyc* and *rheb* in control and *rpS13/+* animals. We found that both *dMyc* and *rheb* over-expression in *rpS13/+* animals rescued timing to the same extent as *rpS13* re-expression, while having minimal effects in the control animals (Figure 5A-B). Recent studies have identified the transcription factor *Xrp1* is an effector of *Minute* phenotypes, particularly cell competition (Ji et al., 2019). However, when we used RNAi to knockdown *Xrp1* specifically in 5-HT neurons we found that there was no effect on developmental timing in the *rpS13/+* animals (Figure 5C).

**Figure 5.**
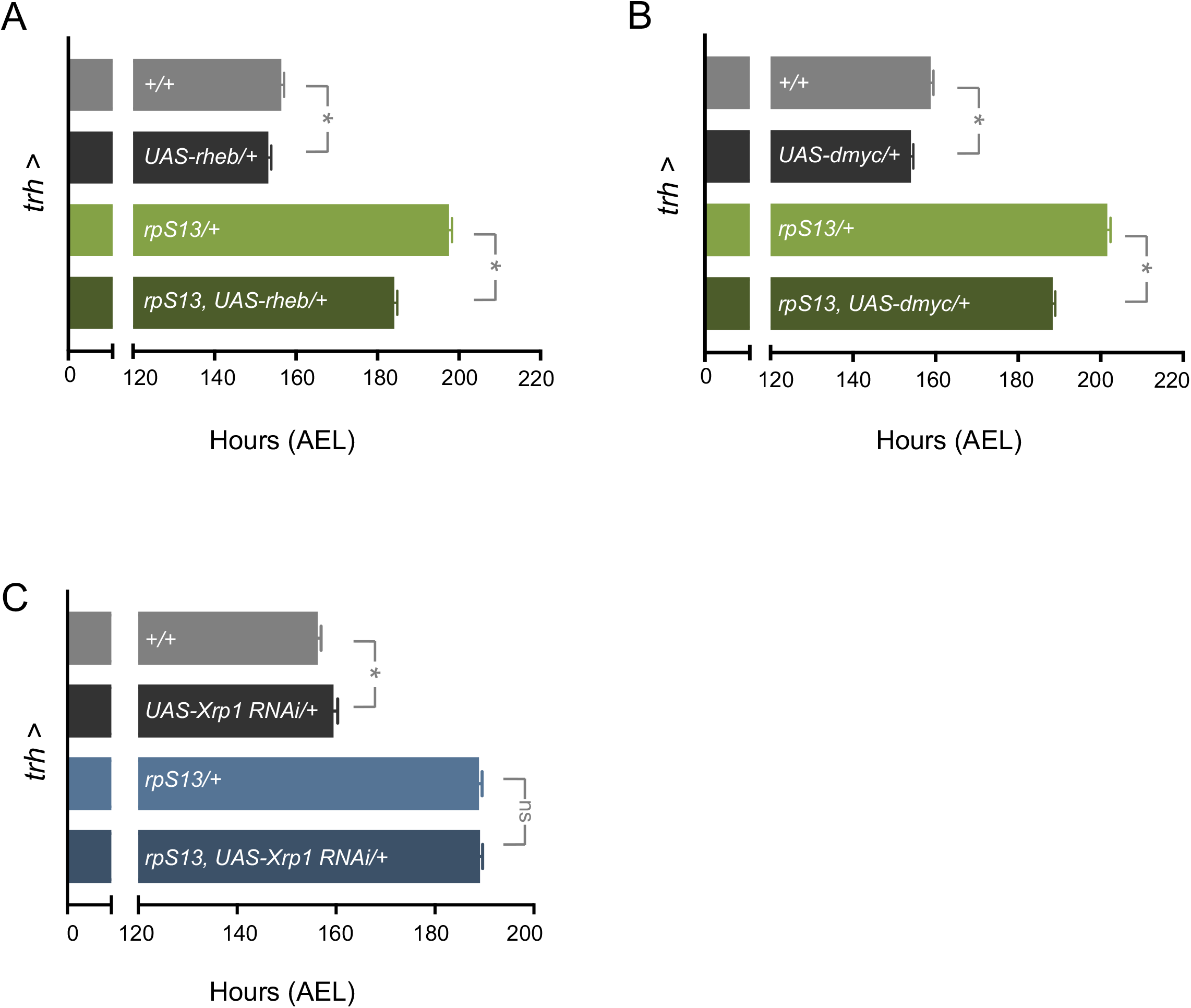
Stimulation of protein synthesis, but not *Xrp1* knockdown in the serotonergic partially rescues *rpS13/+* developmental delay. (A) *dmyc* over-expression in the serotonergic neurons using *Trh-Gal4* partially rescues developmental delay of *rpS13/+* larvae by 13 hours (~31 %). *+/+* n = 198, *UAS-dmyc/+* n = 196, *rpS13/+* n = 173, *rpS13, UAS-dmyc/rpS13* n = 186. Data are presented as +/- SEM. *p < 0.05, Mann-Whitney U test. (B) *rheb* over-expression in the serotonergic neurons using *Trh-Gal4* partially rescues developmental delay of *rpS13/+* larvae by 14 hours (~33 %). *+/+* n = 172, *UAS-rheb/+* n = 235, *rpS13/+* n = 132, *UAS-dmyc/rpS13* n = 185. Data are presented as +/- SEM. *p < 0.05, Mann-Whitney U test. (C) RNAi knockdown of *Xrp1* in serotonergic neurons using *Trh-Gal4* has no effect on developmental timing of *rpS13/+* larvae. *+/+* n = 200, *UAS-Xrp1RNAi/+* n = 160, *rpS13/+* n = 179, *UAS-Xrp1RNAi/rpS13* n = 203. Data are presented as +/- SEM. *p < 0.05, Mann-Whitney U test.

### The developmental delay in *rpS13/+* animals is partially reverse by serotonergic expression of synaptic vesicle proteins

We next focused on what specific aspects of serotonergic neuronal biology might be altered in *rps13/+* larvae to explain their delayed development. It has been previously reported that the 5-HT neurons that project to the PG are regulated by nutrient availability and that in low nutrient conditions these neuronal projections are reduced, leading to diminished 5-HT signaling to the PG and, as a result, reduced ecdysone release and delayed development. We therefore investigated whether *rpS13/+* animals also showed a reduction in axonal projections into the PG. We stained 5-HT neurons in both control and *rpS13/+* animals and in both cases, we observed projections to the PG. When we counted bouton numbers from 5-HT neurons that projected to the PG we found no difference between controls and *rpS13/+* animals suggesting no alterations in 5-HT neuronal outgrowth (Figure 6A-B). It is possible that the 5-HT neurons in *rpS13/+* larvae have reduced activity, leading to decreased stimulation of ecdysone production in the PG. To explore this possibility, we examined the effects of genetic activation of these neurons by expressing the NaChBac sodium channel, which leads to depolarization and activation of neurons. However, we found that expression of *UAS-NaChBac* did not reverse the delay in development seen in *rpS13/+* animals (Figure 6C). We also found that expression of tryptophan hydroxylase (Trh), a key enzyme in the synthesis of 5-HT, also did not alter the delayed development in *rpS13/+* larvae (Figure 6D).

**Figure 6.**
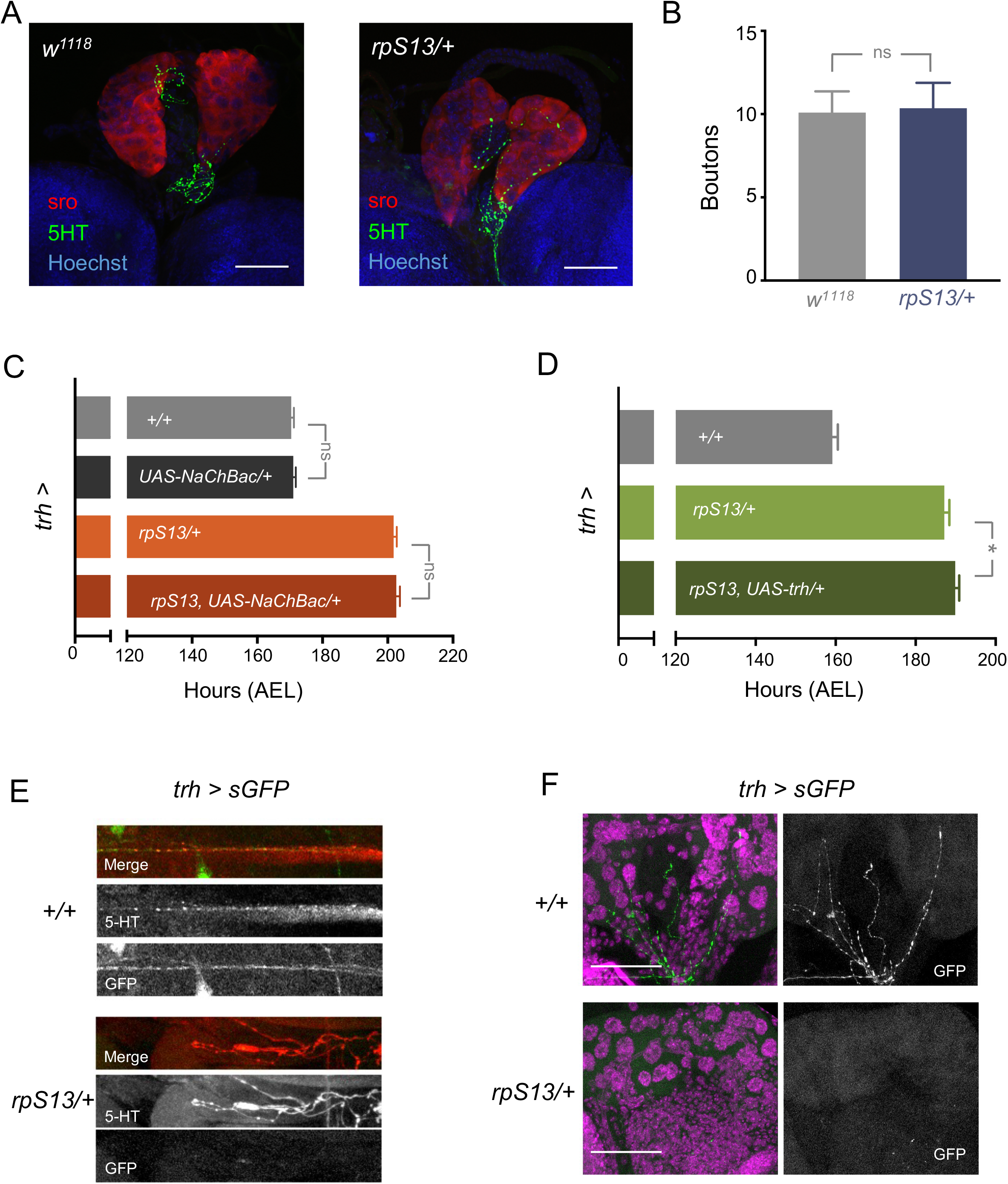
*rpS13/+* animals show normal axonal projections into the prothoracic gland, however are defective in secretory function. (A) Fluorescent confocal images of representative *+/+* and *rpS13/+* prothoracic glands showing innervation of serotonergic neuron termini. Anti-5-HTP is labelled with GFP, while the prothoracic gland is labelled in red using an anti-shroud antibody. Nuclei of both the brain and prothoracic gland are stained with Hoechst. Scale bars, 50μm. (B) Brains of *+/+* (n= 11) and *rpS13/+* (n =11) wandering larvae were dissected and stained for 5-HT. Boutons of serotonin (5-HT) neuron termini that overlap the prothoracic gland were counted. Data are presented as +/- SEM. p > 0.05, Student’s t-test. (C) *UAS-NaChBac* expression in the serotonergic neurons using *trh-Gal4* does not rescue the developmental delay of *rpS13/+* larvae. No difference is seen between *+/+* controls (n = 154) and *UAS-NaChBac* overexpression (n = 135) or between *rpS13/+* (n = 125) and *UAS-NaChBac/rpS13* (n = 108) animals. Data are presented as +/- SEM. p > 0.05, Mann-Whitney U test. (D) *UAS-trh* expression in the serotonergic neurons using *trh-Gal4* does not rescue the developmental delay of *rpS13/+* larvae. *+/+* n = 93, *rpS13/+* n = 71, *UAS-trh/rpS13* n = 91. Data are presented as +/- SEM. *p < 0.05, Mann-Whitney U test. (E) Fluorescent confocal images of representative *trh>sGFP/+* and *trh>sGFP/rpS13* wandering L3 larvae brains. Anti-5-HT is shown in red, while the secreted GFP is green. Images are of 5-HT neurons axons projecting into the PG. (F) Fluorescent confocal images of representative *trh>sGFP/+* and *trh>sGFP/rpS13* prothoracic glands showing innervation of serotonergic neuron termini into the PG. Nuclei of the prothoracic gland are stained with Hoechst (magenta) while secreted GFP from 5-HT neurons are in green. Scale bars, 50μm.

A key process in neurons is the vesicle mediated secretion of neurotransmitters and neuropeptides. We therefore examined whether secretion might be altered in the 5-HT neurons of *rpS13/+* animals. To do this we expressed a secreted form of GFP (sGFP) in the 5-HT neurons of both control and *rpS13/+* animals. In control larvae we saw bright GFP staining in axons and in the termini of neurons that innervate the PG (Figure 6E, F). However, this sGFP expression was strongly suppressed in *rpS13/+* animals (Figure 6E, F) suggesting that they have a secretory defect in their serotonergic neurons (Figure 6F). In contrast with these effects with sGFP, we found that expression of unmodified GFP was similar in controls and *rpS13/+* animals, indicating that the *Minute* animals do not show a general disruption of transgene expression (Figure S3).

Since we saw a defect in the secretory process of *rpS13/+* animals we wanted to further investigate this by looking at proteins required in synaptic secretion. The synthesis, axonal transport, and synaptic release of vesicle contents is controlled by a number of proteins, including members of the SNAP Receptor (SNARE) complex. Interestingly, previous studies have shown that these synaptic vesicle proteins need to be continually synthesized for proper neuronal function (Truckenbrodt et al., 2018) and that their synthesis is often translationally regulated (Batista et al., 2017; Daly and Ziff, 1997). We therefore examined the effects of expressing synaptic vesicle proteins in serotonergic neurons. Strikingly, we found that expression of any of one of three synaptic vesicle proteins - *nsyb, syt1* or *SNAP29* - was able to rescue developmental delay seen in *rpS13/+* animals, to the same magnitude as that seen with expression of *UAS-S13* (Figure 7A-C). We wanted to see if this phenotype was unique to *rpS13/+* animals by examining two other *Minutes, rpS24/+* and *rpS26/+*. Like *rpS13/+*, these two additional *Minutes* also show a delay in development, albeit slightly weaker. However, as with *rpS13/+* animals, we saw that this delay in timing was also rescued by about a third, by overexpression of *nsyb* (Figure 7D-E).

**Figure 7.**
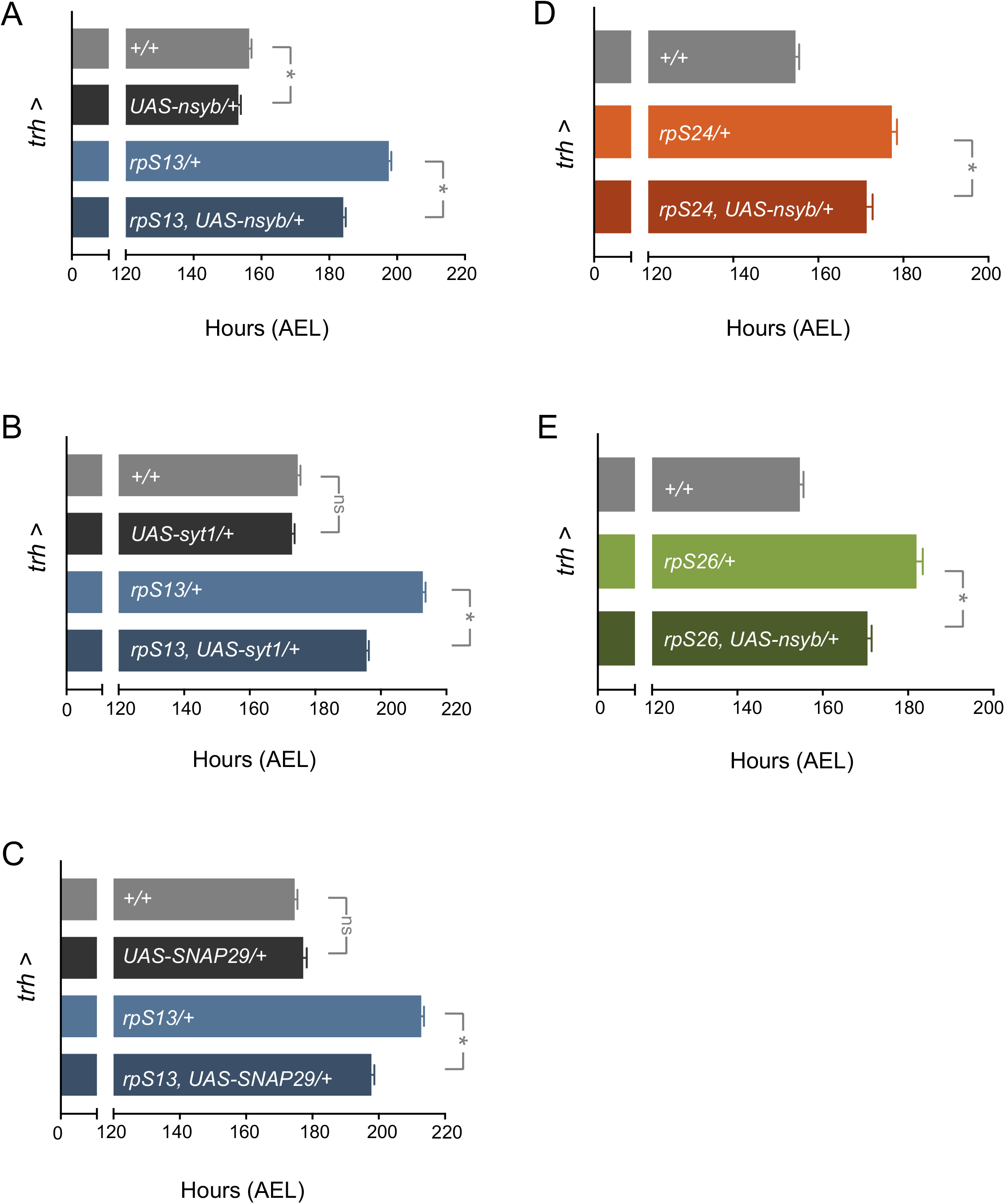
Expression of individual SNARE complex proteins partially rescues *rpS13/+* developmental delay as well as the other *rpS24/+* and *rpS26/+* developmental delay. (A) *nsyb* over-expression in the serotonergic neurons using *Trh-Gal4* partially rescues developmental delay of *rpS13/+* larvae by 12 hours (~30 %). *+/+* n = 182, *UAS-nsyb/+* n = 180, *rpS13/+* n = 190, *rpS13, UAS-nsyb/rpS13* n = 176. Data are presented as +/- SEM. *p < 0.05, Mann-Whitney U test. (B) *syt1* over-expression in the serotonergic neurons using *Trh-Gal4* partially rescues developmental delay of *rpS13/+* larvae by 17 hours (~43 %). *+/+* n = 168, *UAS-syt1/+* n = 151, *rpS13/+* n = 137, *rpS13, UAS-syt1/rpS13* n = 136. Data are presented as +/- SEM. *p < 0.05, Mann-Whitney U test. (C) *SNAP29* over-expression in the serotonergic neurons using *Trh-Gal4* partially rescues developmental delay of *rpS13/+* larvae by 15 hours (~39 %). *+/+* n = 168, *UAS-SNAP29/+* n = 140, *rpS13/+* n = 137, *rpS13, UAS-SNAP29/rpS13* n = 135. Data are presented as +/- SEM. *p < 0.05, Mann-Whitney U test. (D) *nsyb* over-expression in the serotonergic neurons using *Trh-Gal4* partially rescues developmental delay of *rpS24/+* larvae by 6 hours (~26 %). *+/+* n = 168, *rpS24/+* n = 105, *UAS-nsyb/rpS24* n = 124. Data are presented as +/- SEM. *p < 0.05, Mann-Whitney U test. (E) *nsyb* over-expression in the serotonergic neurons using *Trh-Gal4* partially rescues developmental delay of *rpS26/+* larvae by 11.5 hours (~42 %). *+/+* n = 168, *rpS13/+* n = 98, *rpS26, UAS-nsyb/rpS26* n = 120. Data are presented as +/- SEM. *p < 0.05, Mann-Whitney U test.

## DISCUSSION

Since the discovery that *Minutes* are mutants for Rps, a prevailing hypothesis (the “balance hypothesis”) has been that their developmental phenotypes result from an imbalance of cytoplasmic ribosomal protein concentrations leading to incomplete ribosomal subunit assembly and reduced overall translation (Marygold et al., 2007). However, our data indicate that despite all cells in a *Minute* animal being heterozygous for an Rp gene, this does not result in a whole-body decrease in ribosome numbers or protein synthesis. Rather, the developmental delay phenotypes in *Minute* animals likely result from more selective effects of Rp loss. Here we identify 5-HT neurons as one important cell-type in which Rp function is required for proper larval developmental timing.

Interestingly we found that the effect of RpS13 in 5-HT neurons could account for 30-40% of the total developmental delay in *rps13/+* animals. This suggests that RpS13 containing ribosomes may be playing roles in other tissues to control developmental timing. Indeed, previous studies have also shown tissue-selective effects of Rps on development. For example, re-expression of *rpS6* in prothoracic glands could partially rescue the animal’s developmental delay in *rpS6/+* larvae (Lin et al., 2011), although we did not find such a PG specific role for *rpS13*. It has also been shown that disc-specific knockdown of dilp8 in *rpS3/+* animals can partially reverse their larval developmental delay (Akai et al., 2021). One possibility, therefore, is that Rp function in a combination of tissues is required for proper endocrine control of developmental timing. Further studies are required to see if these tissue-specific contributions are similar for all Rps or may show some heterogeneity depending on the Rp being examine. The 5-HT neurons that innervate the PG have been shown to be particularly important in coupling nutrition to the control of ecdysone and developmental timing (Shimada-Niwa and Niwa, 2014). Given that the control of ribosome synthesis is a conserved function of nutrient-signaling pathways, our results pinpointing a key role for Rps in the function of these neurons suggest one way that nutrients may modulate the 5-HT control of ecdysone.

Our work suggests that the requirement for RpS13 in 5-HT neurons may reflect a role in the control of vesicle-mediated secretion. We saw that expression of a secreted form of GFP was lost on the axons and termini of 5-HT neurons, and we saw that expressing different vesicle proteins in these neurons in *rps13/+* animals was able to rescue the developmental delay to the same extent as *UAS-rpS13* re-expression. How might loss of S13 affect secretion in 5-HT neurons? One plausible mechanism is through reduced translation. This notion is supported by our results showing that serotonergic expression of two conserved inducers od protein synthesis, Myc and Rheb, could rescue the delay in development seen in *rp13/+* animals. Current models suggest that rp/+ phenotypes arise from either heterogeneous ribosomes or selective mRNA translational control (Dinman, 2016; Genuth and Barna, 2018a, b; Khajuria et al., 2018; Mills and Green, 2017). We saw a partial rescue in timing with *UAS-nsyb* overexpression in three different *minutes - rps13/+, rps24/+* and *rpS26/+* - suggesting perhaps that ribosome heterogeneity may not explain the 5-HT neuronal contribution to the *Minute* phenotypes. Alternatively, a reduction of RpS13 in these neurons may result in selective changes in mRNA translation of proteins involved in controlling neuronal secretory function. In mammalian systems, synaptic vesicle proteins are required to be continually synthesized for proper neuronal function (Truckenbrodt et al., 2018). Moreover, their expression has been shown to be post-transcriptionally regulated (Daly and Ziff, 1997). Hence, the vesicle proteins themselves may be subject to selective translational regulation. It could be that SNARE complex mRNAs share a common feature in their 5’ or 3’ UTR regions that leads to them being translationally regulated in similar ways. Indeed, UTR-mediated regulation of translation is a prevalent mode of controlling gene expression in neurons (Blair et al., 2017). The subcellular control of translation is also important in neurons (Biever et al., 2019; Cioni et al., 2018; Holt et al., 2019). Here, local translation in regions such as cell bodies, axons and termini allows for selective, spatial control over protein synthesis. For example, it has been found that SNAP25 synaptic vesicle proteins can be locally translated in axons (Batista et al., 2017). Thus, it is possible that the defects in *Minute* animals may occur due to alterations in local translation in 5-HT neurons.

Our findings may also be relevant to human biology. It has been observed that many neurological disorders arise from abnormal synaptic vesicle formation and function. These “synaptopathies” include disorders such as schizophrenia, AHDH, and autism, and are often associated with aberrant mRNA translation (Bagni and Zukin, 2019; Chen et al., 2019). Interestingly some ribosomopathies in humans have also been shown to present with various neurological disorders such as microcephaly and mental retardation. In particular, mutations in *rpS23* and *rpL10* have been shown to be associated with autism spectrum disorder (Klauck et al., 2006; Paolini et al., 2017). Perhaps certain ribosomopathies that present mental disorders could in part be due to defect in vesicle function and, as a result, disrupted synaptic function.

## MATERIALS AND METHODS

### Fly strains and husbandry

Flies were raised on food with the following composition: 150 g agar, 1600 g cornmeal, 770 g torula yeast, 675 g sucrose, 2340 g D-glucose, 240 ml acid mixture (propionic acid/phosphoric acid per 34 liters of water). For all experiments larvae were maintained at 25°C, unless otherwise indicated. Unless otherwise stated the fly strains used were obtained from the Bloomington Stock Center (BDSC): *w^1118^, yw, yw*;P[w[+mC]=lacW]rpS13[1]/cyoGFP* (2246)*, UAS-rpS13GFP/cyo* (Gift from S. Brogna)*, P0206-GAL4* (Gift from C. Mirth)*, esg^ts^-GAL4, elav-GAL4* (Gift from F.Buldoc)*, nsyb-GAL4* (51635)*, ptth-GAL4* (Gift from M. O’Connor), *trh-GAL4* (38388)*, GMR29H01-GAL4* (47343)*, UAS-dmyc-dp110* (25914)*, UAS-rheb* (9689)*, UAS-NaChBac* (9469)*, UAS-trh* (27638)*, UAS-nsybGFP* (6921, 6922)*, UAS-syt1* (6925)*, UAS-SNAP29* (56817)*, P0206-GAL4* (Gift from M. O’Connor), *lgr3-GAL4* (66683)*, UAS-Xrp1RNAi* (34521)*, w ^1118^; PBac[w[+m]]=WH] rpS24[f06717]/cyoGFP* (19002)*, P[ry[+t7.2]=PZ]rpS26[04553]/cyoGFP(12048), UAS-sGFP* (Gift from M. Gonzlez-Gaitan).

### Measurement of *Drosophila* development and body size

For measuring development timing to pupal stage, newly hatched larvae were collected at 24 hr AEL and placed in food vials (50 larvae per vial) and kept at 25°C. The number of pupae was counted twice a day. For each experimental condition, a minimum of four replicates was used to calculate the mean time to develop into pupae. To measure pupal volume, pupae were imaged using a Zeiss Discovery.V8 Stereomicroscope with Axiovision imaging software. Pupal length and width were measured, and pupal volume was calculated using the formula, volume = 4/3π(L/2) (l/2)2. A minimum of four replicates was used to calculate the mean volume for each genotype.

### Quantitative PCR

Total RNA was extracted from larvae (groups of 10) using TRIzol according to manufacturer’s instructions (Invitrogen; 15596-018). RNA samples were then subjected to DNase treatment according to manufacturer’s instructions (Ambion; 2238 G) and reverse transcribed using Superscript II (Invitrogen; 100004925). The generated cDNA was used as a template to perform qRT–PCRs (ABI 7500 real time PCR system using SyBr Green PCR mix) using specific primer pairs. PCR data were normalized to either actin or alpha-tubulin levels. The following primers were used:

RpS13 forward: AGGCAGTGCTCGACTCGTAT
RpS13 reverse: TTCCCGAGGATCTGTACCAC

Beta-tubulin forward: ATCATCACACACGGACAGG
Beta-tubulin reverse: GAGCTGGATGATGGGGAGTA

Actin5C forward: GAGCGCGGTTACTCTTTCAC
Actin5C reverse: GCCATCTCCTGCTCAAAGTC

18S rRNA forward: CCTGCGGCTTAATTTGACTC
18S rRNA reverse: ATGCACCACCACCCATAGAT

28S rRNA forward: TGCCAGGTAGGGAGTTTGAC
28S rRNA reverse: CAAGTCAGCATTTGCCCTTT

spookier forward: TATCTCTTGGGCACACTCGCTG
spookier reverse: GCCGAGCTAAATTTCTCCGCTT

phantom forward: GGATTTCTTTCGGCGCGATGTG
phantom reverse: TGCCTCAGTATCGAAAAGCCGT

### Puromycin assay

Groups of 10 wandering larvae or earlier time point larvae were inverted in Schneider’s media and then transferred to Eppendorf tubes containing media plus 5 μg/ml puromycin (Sigma), 6 different replicates were used for Figure 1F. The larval samples were then left to incubate on a mutator for 40 minutes at room temperature. Following incubation, the inverted larvae were snap frozen for western blot analyses. For experiments on larval brains, inverted larvae were placed in ice-cold PBS after incubation with puromycin, and the brains were isolated and lysed for western blot analyses.

### Western Blotting

Whole inverted larvae or isolated brains were lysed with a buffer containing 20 mM Tris-HCl (pH 8.0), 137 mM NaCl, 1 mM EDTA, 25% glycerol, 1% NP-40 and with following inhibitors: 50 mM NaF, 1 mM PMSF, 1 mM DTT, 5 mM sodium ortho vanadate (Na3VO4) and protease inhibitor cocktail (Roche cat. no. 04693124001) and phosphatase inhibitor (Roche cat. no. 04906845001), according to the manufacturer’s instruction. Protein concentrations were measured using the Bio-Rad Dc Protein Assay kit II (5000112). For each experiment, equal amounts of protein lysates for each sample (40 μg) were resolved by SDS-PAGE and electrotransferred to a nitrocellulose membrane. Blots were then briefly stained with Ponceau S to visualize total protein and then subjected to western blot analysis with specific antibodies. Protein bands were then visualized by chemiluminescence (enhanced ECL solution, Perkin Elmer). Primary antibodies used were anti-puromycin (3RH11) antibody (1:1000, Kerafast, Boston, USA, cat. no. EQ0001), anti-eIF2alpha (1:1000, AbCam #26197). Secondary antibodies were purchased from Santa Cruz 144 Biotechnology (sc-2030, 2005, 2020, 1: 10,000).

### 20-Hydroxyecdysone Feeding

Newly hatched *w^1118^* and *rpS13/+* larvae were collected at 24 hr AEL and placed in food vials (50 larvae per vial) supplemented either with 20-hydroxyecdysone (Sigma-Aldrich CAS number 5289-74-7) or equal volume of 95% ethanol for controls. 20-hydroxyecdysone was dissolved in 95% ethanol to a final concentration of 0.3mg/ml. A minimum of four replicates was used to calculate the mean volume for each genotype.

### Immunostaining and microscopy

Drosophila larvae were fixed in 8% paraformaldehyde/PBS at room temperature for 30 min. After blocking for 2 h in 1% BSA in PBS/0.1% Triton-X 100, inverted larvae were incubated overnight in anti-5HT (Sigma-Aldrich AB125) and anti-shroud antibody (gift from R.Niwa) (1:2000, 1:500 dilutions). Primary antibody staining was detected using Alexa Fluor 488 (Molecular Probes) goat-anti rabbit secondary antibodies. Brains with attached prothoracic glands were then dissected out and mounted on coverslips using mounting media (Vectashield).

### Bouton quantification

Brain and attached prothoracic glands were dissected from *w^1118^* and *rpS13/+* larvae during the wandering L3 stage and were stained with anti-5HT and anti-shroud antibodies. Confocal Z stack images were acquired and 5-HT boutons that overlapped the PG (based on anti-shroud staining) were counted. The values for each individual brain were recorded and averages for each genotype were calculated.

### Statistics

For all experiments, error bars represent standard error of mean (SEM). Data were analyzed by Students t-test or Mann-Whitney U test. All statistical analysis and data plots were performed using Prism software. In all figures, statistically significant differences are presented as * and indicate p<0.05.

## Supplemental Figures

**Figure S1.**
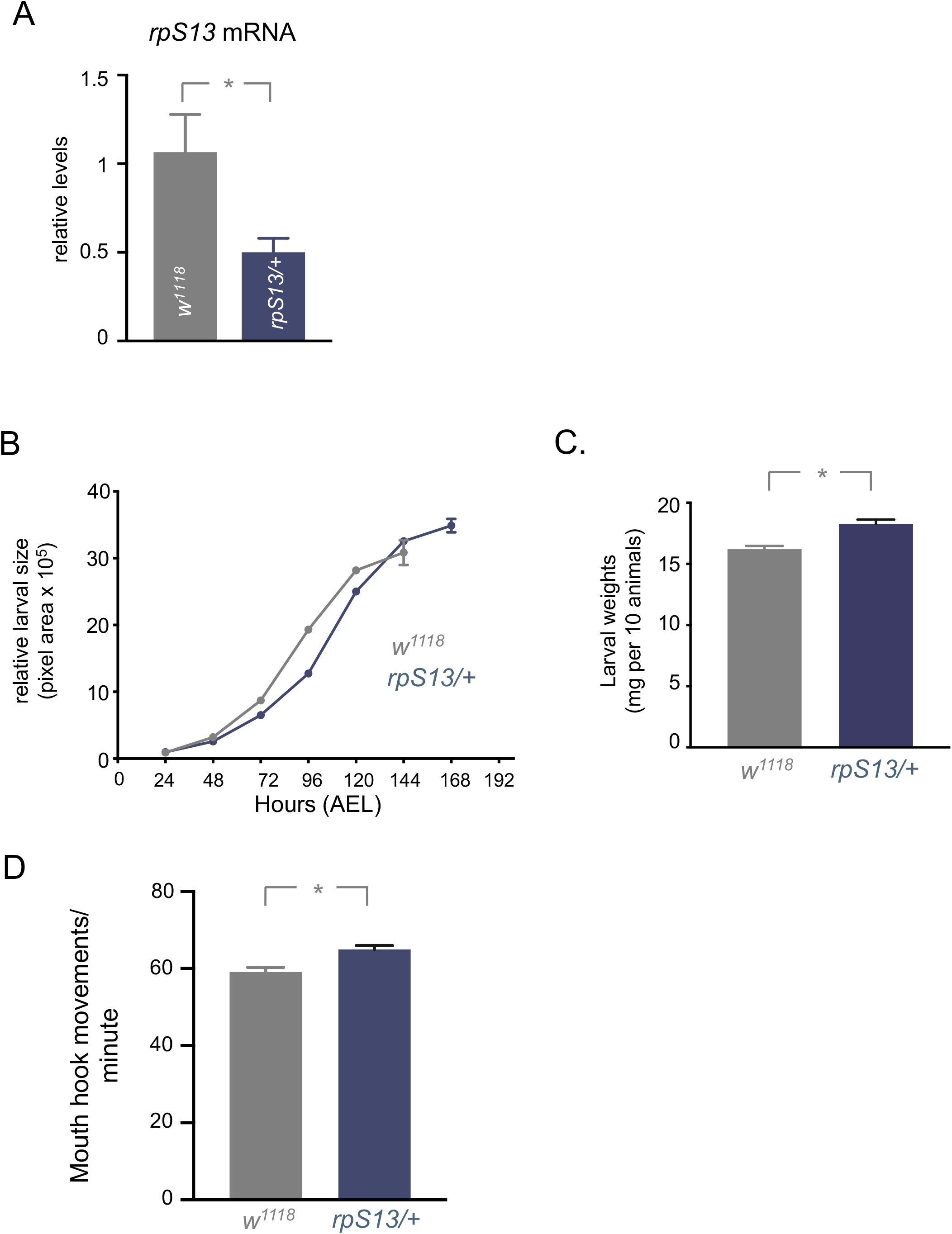
(A). Transcript levels of the *rpS13* heterozygotes are reduced to half the wild-type levels present in controls. mRNA was isolated from third instar wandering larvae. Total RNA was isolated and measured by qRT-PCR, n = 4 independent samples per genotype. Data are presented as +/- SEM. *p < 0.05, Student’s t-test. (B) Relative larval size of *W^1118^* and *rpS13/+* animals throughout development. Larval area was measured every 24 hours after hatching until wandering and recorded as pixel area. Data are presented as +/- SEM. (C) Larval weight (mg) at wandering L3 stage of *W^1118^* and *rpS13/+* animals. Larvae were measured in groups of 10, n = 10 independent samples per genotype. Data are presented as +/- SEM. *p < 0.05, Student’s t-test. (D) Mouth hook movements recorded in one minute of feeding for 96-hour L3 larvae of *W^1118^* and *rpS13/+* animals. *W^1118^* n = 20, *rpS13/+* n = 20. Data are presented as +/- SEM. *p < 0.05, Student’s t-test.

**Figure S2.**
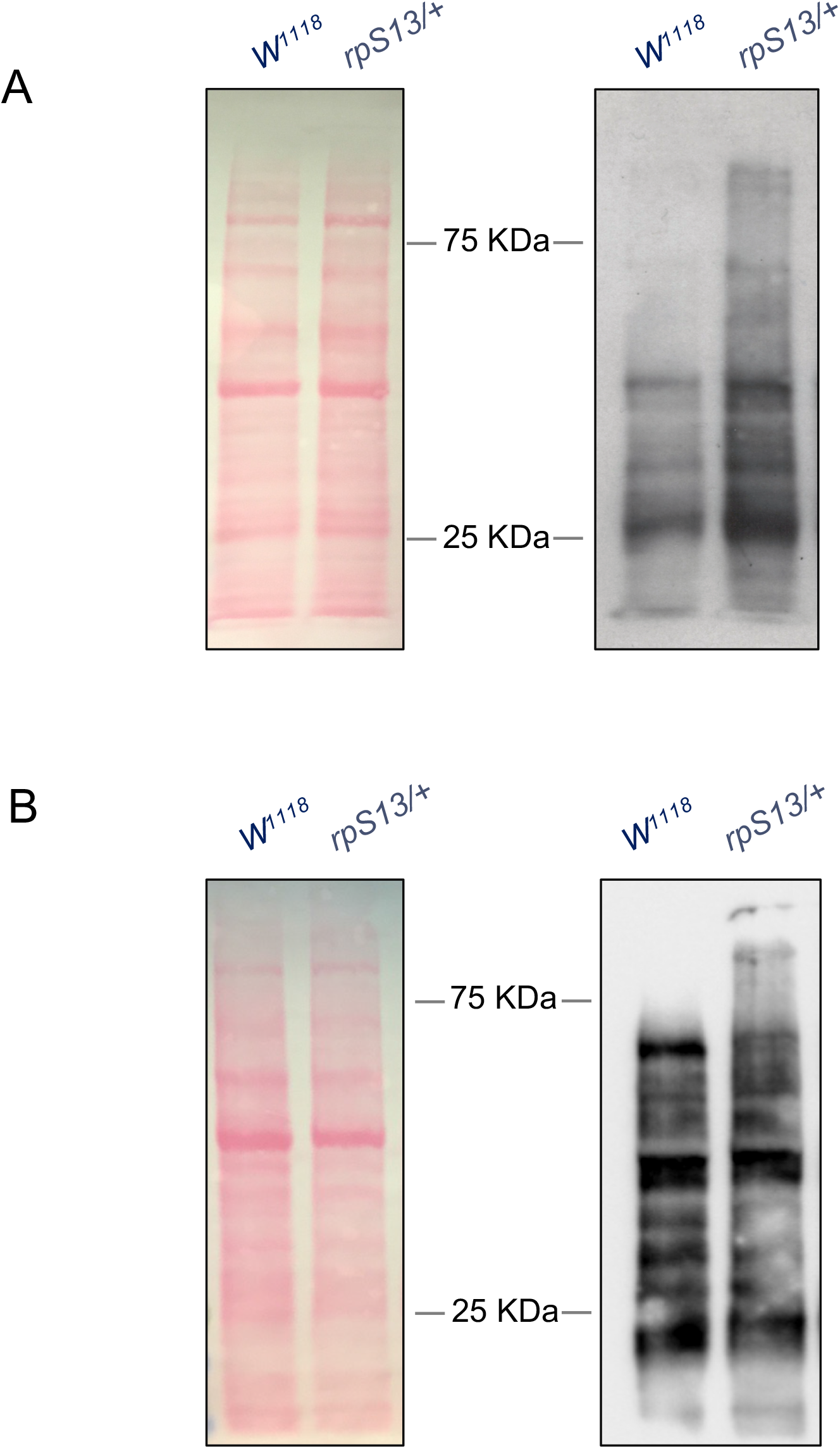
Puromycin labelling assay shows no decrease, but rather a small increase in *rpS13/+* translational rates in whole larvae when compared to age matched controls. (A) 96-hour *W^1118^* and *rpS13/+* larvae. Left, Ponceau S staining showing total protein. Right, anti - puromycin immunoblot. (B) 120-hour *W^1118^* and *rpS13/+* larvae. Left, Ponceau S staining showing total protein. Right, anti - puromycin immunoblot.

**Figure S3.**
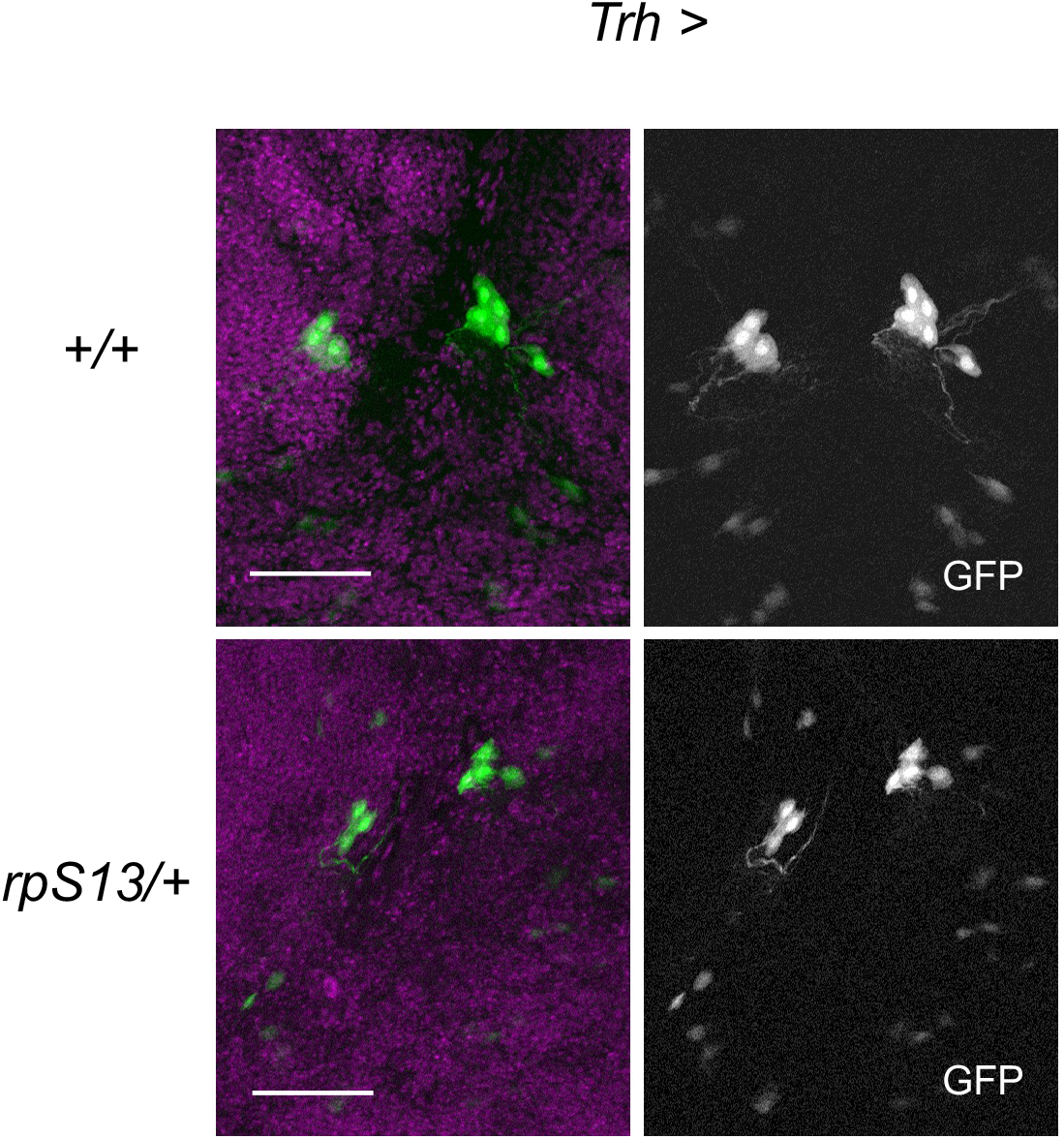
Fluorescent confocal images of representative *trh>GFP/+* and *trh>GFP/rpS13* wandering L3 larval brains. Nuclei of neurons stained with Hoechst (magenta) while 5-HT neurons are labelled with GFP. Scale bars, 50μm.

